# Left-to-right dorsomedial prefrontal cortex interhemispheric projections mediate psychosocial stress vulnerability

**DOI:** 10.1101/2025.08.07.669148

**Authors:** Gessynger Morais-Silva, Beatriz Fagundes Gasques, Isabelle Lima Lugli, Ricardo Luiz Nunes-de-Souza

**Author notes:** **Corresponding author:** Current address: Department of Pharmacology and Therapeutics, State University of Maringá, 87020-900 Maringá, PR, Brazil, E-mail address (G. Morais-Silva).

## Abstract

Functional asymmetries in the medial prefrontal cortex (mPFC) are significant attributes of this brain area, implicated in its role in emotional processing and executive function. Evidence suggests that, under normal conditions, there is a tonic inhibition of the right (R) mPFC by the left (L) mPFC, and a dysregulation of this hemispheric functional lateralization is implicated in detrimental chronic stress effects. Considering the wide interhemispheric connection and the inhibitory tone from the LmPFC to the RmPFC, we hypothesize that alterations in the activity of the direct projections between the mPFC hemispheres during stressful situations are related to stress vulnerability. To address this question, we used a chemogenetic approach to modulate the activity of L-to-R dorsomedial prefrontal cortex (dmPFC) monosynaptic projections during psychosocial stress (PSS) exposure in mice. We found that activating LdmPFC projections during a repeated PSS protocol prevents stress-induced apathy-like behavior in females and males and social avoidance in male mice. On the other hand, inhibiting such projections during a single session of PSS increases vulnerability to stress effects in male mice, increasing social avoidance and anxiety-like behaviors. Both glutamatergic and GABAergic cells compose the projecting interhemispheric neurons in the dmPFC. However, the LdmPFC showed a higher density of glutamatergic projections to the RdmPFC than the opposite. In conclusion, our results revealed an involvement of the monosynaptic projections from the LdmPFC to RdmPFC in the vulnerability to the behavioral alterations induced by PSS in female and male mice.

## 1. Introduction

Functional hemispheric asymmetries, or functional brain lateralization, are a fundamental feature of cerebral structure, impacting from motor functions to cognitive processing (Ocklenburg and Güntürkün, 2012). Not surprisingly, many psychiatric disorders have been linked to alterations in functional brain lateralization (Ocklenburg et al., 2024). In this context, the extensive communication between the medial prefrontal cortex (mPFC) hemispheres, especially its dorsal part (dmPFC) (Hoover and Vertes, 2007) and seminal findings from the 1970s to late 1990s (Gainotti, 1972; Robinson et al., 1984; Robinson and Szetela, 1981; Sullivan and Gratton, 2002, 1999) have led to the hypotheses that a disruption in the functional asymmetries in this region underlies the development of stress-related disorders. Central to this framework is the postulate that, under baseline conditions, there is a tonic inhibition of the right (R) dmPFC by the left (L) dmPFC. However, chronic stress exposure disrupts this inhibitory control, heightening RdmPFC activity to drive emotional expression, promoting anxiogenic and depressogenic responses, and reducing behavioral flexibility (Cerqueira et al., 2008).

Consistent findings across human and animal research support this hypothesis. Patients affected by ischemic lesions in the anterior part of the left frontal cortex show an increase in anxious/depressive mood compared to those affected by lesions in any other brain sites (Gainotti, 1972; Robinson et al., 1984; Robinson and Szetela, 1981). Moreover, the severity of the symptoms is correlated to the proximity of the lesion to the frontal pole in the computed tomography scan (Robinson et al., 1984). Lower left frontal cortex baseline activity, relative to the right frontal cortex, in women, is correlated to an increase in the severity of depressive symptoms in the future (Stewart and Allen, 2018). In male rats, ibotenic acid-induced lesions of the RmPFC attenuate neuroendocrine responses to repeated restraint stress (RRS) (Sullivan and Gratton, 1999) and decrease anxiety-like behaviors in the elevated plus maze (EPM) (Sullivan and Gratton, 2002). Sustained hypercorticosteronemia in male rats decreases left anterior cingulate cortex (ACC) volume (Cerqueira et al., 2005), while aged men that do not show a decrease in plasmatic cortisol concentration after a low-dose dexamethasone administration (called non-suppressors) have smaller left ACC volume relative to suppressors, i.e., those who show a decrease in plasmatic cortisol after low-dose dexamethasone administration (MacLullich et al., 2006). The intrinsic differences in dendritic arborization and neurogenesis between the LdmPFC and the RdmPFC in male rats are abolished following chronic social defeat stress (CSDS) (Czéh et al., 2007) or RRS exposure (Perez-Cruz et al., 2007).

While few studies have assessed the effects of unilateral modulation of the dmPFC activity, growing evidence corroborates the hypothesis of dmPFC functional lateralization in mediating stress-related outcomes. The optogenetic activation of the LmPFC reversed the social avoidance induced by CSDS exposure in male mice, while the optogenetic inhibition of the LmPFC induced social avoidance in individuals identified as resilient to the CSDS (Lee et al., 2015). The nitrergic activation of the RdmPFC using a nitric oxide donor produces anxiogenic-like effects, whereas RdmPFC inactivation using cobalt chloride elicits anxiolysis in male mice exposed to the EPM (Costa et al., 2016). Male mice subjected to the CSDS showed an increased activation of the nitrous oxide-producing neurons in the RmPFC (Faria et al., 2020). The inactivation of the LdmPFC using cobalt chloride prolonged the anxiogenic-like effect of a single social defeat stress (SDS) episode (Costa et al., 2016; Victoriano et al., 2020). Additionally, CSDS exposure increases the activation of glutamatergic neurons specifically in the right prelimbic area (PrL) (Santos-Costa et al., 2021). In a model of post-traumatic stress disorder (PTSD) in male rats, daily optogenetic activation of the left infralimbic cortex (IL) rescues fear extinction deficits (Canto-de-Souza et al., 2021), while RmPFC inactivation using cobalt chloride reduces the conditioned freezing behavior but not cardiovascular responses in contextual fear conditioning in male rats (Gomes-de-Souza et al., 2024).

Collectively, the evidence above indicates that the functional asymmetry and interhemispheric communication within the dmPFC play a critical role in emotional responses to stressful stimuli, and their disruption may impact how individuals cope with stressful environments and adverse situations. However, it is not clear yet whether a disruption in the activity of the LdmPFC interhemispheric projections is related to the abovementioned dysregulation of the dmPFC functional lateralization induced by chronic stressors. Here, we examined the impact of the LdmPFC activity on the behavioral consequences of psychosocial stress (PSS) exposure.

## 2. Material and methods

### 2.1. Subjects

One hundred and forty-three Swiss-Webster mice (70 males and 73 females, 6-8 weeks old on the surgery day) were used as experimental subjects. Additionally, 20 male Swiss-Webster mice (>4 months old) were used as aggressors in the PSS. Animals were obtained from the *Centro de Pesquisa e Produção de Animais* (CPPA) at the São Paulo State University/UNESP (Botucatu, Brazil) and arrived at the local animal facility at 3 weeks old. Experimental subjects were grouped (age and sex-matched, 4-5 animals/cage) in individually ventilated polysulfone cages (240 x 251 x 386 mm, floor area 695 cm^2^, ref. 000121, ALBR Indústria e Comércio LTDA, Monte Mor, BRA) with sawdust bedding supplied with sterile cardboard tubes as environmental refinement (Ambient Trio®, Animal Pro, São Paulo, BRA). The facility was maintained under an artificial 12h light-dark cycle (lights on at 7 am), and controlled temperature (23 ± 1°C). Tap water (supplied by the distribution network) and food (Benelab, Qualy Nutrição Animal Indústria e Comércio Ltda., Lindóia, BRA) were freely available except during the brief periods of behavioral testing. The aggressors used in the PSS were individually housed in individually ventilated polysulfone cages (207 x 216 x 316 mm, floor area 451 cm^2^, ref. 003466, ALBR) under the same conditions as described above.

All procedures involving animals were previously approved by the local ethics committee for animal care and use (CEUA FCF/CAr protocol number 16/2021) and were conducted according to the principles of the Brazilian Guide for the Care and Use of Animals in Scientific Research (DBCA) issued by the National Council for Animal Experiments Control (CONCEA/MCTI). The DBCA was based on the NIH (National Research Council) Guide for the Care and Use of Laboratory Animals.

### 2.2. Clozapine N-oxide, Fluoro-Gold, and viral vectors

Clozapine-N-oxide base (CNO) was obtained from the National Institute of Mental Health (NIMH) Chemical Synthesis and Drug Supply Program (catalog number C-929). It was dissolved (0.1 mg/mL) in a vehicle solution containing dimethyl sulfoxide (DMSO, 1%) and 0.1M phosphate-buffered saline (PBS). CNO was used to activate the designer receptors exclusively activated by designer drugs (DREADDs) and was administered in a dose of 1 mg/kg (0.1 ml/10 g, i.p.) based on our previous publication (Morais-Silva et al., 2023). An experiment was conducted to evaluate whether the CNO dose used in our study can induce behavioral alterations on its own. The full experimental description and the obtained data are available as supplementary material (Supplementary Material S1, Figure S1).

For the manipulation of the monosynaptic dmPFC interhemispheric projections, we used a combination of the following adeno-associated viral vectors (AAV): pENN.AAV5.hSyn.HI.eGFP-Cre.WPRE.SV4 (Addgene viral prep #105540-AAV5, a gift from James M. Wilson), hereafter abbreviated as AAV5-Cre; rgAAV-hSyn-DIO-hM3Dq-mCherry (Addgene viral prep #44361-AAVrg, a gift from Bryan Roth), hereafter abbreviated as rgAAV-DIO-hM3Dq; rgAAV.hSyn.DIO.hM4Di.mCherry (Addgene viral prep #44361-AAVrg, a gift from Bryan Roth) hereafter abbreviated as rgAAV-DIO-hM4Di; rgAAV-hSyn-DIO-mCherry (Addgene viral prep #50459-AAVrg, a gift from Bryan Roth) hereafter abbreviated as rgAAV-DIO-mCherry;

The neuronal retrograde tracer Fluoro-Gold^TM^ (Fluorochrome LLC, Englewood, USA), dissolved at a concentration of 1% in saline (NaCl 0.9%) was used for the characterization of the monosynaptic dmPFC interhemispheric projections.

### 2.3. Stereotaxic surgery

Experimental animals were subjected to a stereotaxic surgery for the infusion of 0.1 µL of the desired AAV or Fluoro-Gold^TM^ in the LdmPFC or RdmPFC depending on the experiment. For that, mice were anesthetized with isoflurane (5% induction, 3-1.5% maintenance in 21% oxygen at 0.8 L/min, Isoforine®, Cristália, Itapira, BRA) using a digital low-flow inhalation anesthesia system (Bonther, Ribeirão Preto, BRA), had their head fixed on the stereotaxic frame (51615T, Stoelting Co., Wood Dale, USA) for targeting the LdmPFC or the RdmPFC (anteroposterior, 1.9 mm; mediolateral, 0.3 mm to the left or the right hemisphere; dorsoventral, −2.0 mm from the skull surface, 90° angled). The microinjection was performed using a microsyringe (0.5 µL Neuros Syringe, Model 7000.5 Knurled Hub, 32G, Hamilton Co., Reno, USA) coupled with a precision pump (Stoelting Quintessential Stereotaxic Injector, Stoelting Co.) at 0.02 µL/min. After the infusion, the needle was kept in place for an additional 5 min to avoid liquid diffusion. After the microinjection, the skin was sutured (5-0 Nylon suture, Labor Import Comercial Imp. e Exp. Ltda, BRA) and the animals were monitored until the beginning of the experiments. Right before the surgery, animals received a prophylactic administration of a veterinary antibiotic (17 mg/mL; 0.1 mL/animal i.m., Pentabiótico® Veterinário, Zoetis, BRA; a commercial mixture of 600,000 UI benzathine benzylpenicillin; 600,000 UI procaine benzylpenicillin; 300,000 UI potassium benzylpenicillin; 250 mg dihydrostreptomycin; 250 mg streptomycin) and the nonsteroidal anti-inflammatory drug flunixin-meglumine (3.5 mg/kg s.c., Banamine®, Merck & Co., Inc., Rahway, USA). The animals were allowed to recover for at least 2 weeks before the beginning of the experiments.

### 2.4. Histological determination of microinjection site and Immunofluorescence

At the end of the experiments, mice were deeply anesthetized with urethane (2.5 g/kg, 0.1 mL/10 g i.p., U2500, Sigma-Aldrich Co., St. Louis, USA), and transcardially perfused with 24 mL of room temperature 0.1M PBS pH 7.4 followed by 50 mL of cold (4°C) 4% paraformaldehyde (PFA). The brains were removed, kept in PFA at 4°C for 24 hours, and then transferred to a 30% sucrose solution in PBS for 48 hours at 4°C. Then, the brains were flash-frozen and stored at −80°C until needed. Coronal sections of the dmPFC (30 µm) were obtained in PBS using a cryostat (Leica CM1850, Leica Biosystems, Nussloch, GER). To the correct identification of the brain hemisphere, the right hemisphere was marked using a 25G blunt needle before sectioning.

For the histological determination of the microinjection site in experiments 1 and 2, slices were washed in PBS (3x 10-minute washes), mounted in adhesion microscope slides (158105B, Indústria e Comércio de Produtos Científicos Perfecta LTDA, São Paulo, BRA) with Fluoroshield™ mounting media (F6182, Sigma-Aldrich Co.) and sealed with coverslip and nail polish. Images of the LdmPFC and RdmPFC were captured at 25x and 200x magnification on an epifluorescence microscope (Axio Imager.D2, Carl Zeiss Microscopy, LLC, Thornwood, USA) using the Zen Pro 2.0 software (Carl Zeiss Microscopy) for the visualization of eGFP and mCherry fluorescence. Animals showing eGFP or mCherry fluorescence in brain areas other than the dmPFC, no expression of the reporter fluorescent proteins, or bilateral eGFP expression (result of AAV5-Cre spillage) were excluded from the analysis.

For the characterization of the interhemispheric projections in experiment 3, slices were washed in PBS (3x 10-minute washes) and blocked in a solution containing 5% horse serum (H1270, Sigma-Aldrich Co.), 3% bovine serum albumin (Interlab LTDA, São Paulo, BRA) and 0.2% Triton X-100 (Dinâmica Química Contemporânea LTDA, Indaiatuba, BRA) in PBS for 1 hour. Then, slices were washed in PBS (6x 5-minute washes) and then incubated overnight at 4°C in blocking buffer containing the primary monoclonal antibodies mouse anti-CAMKII alpha (1:200, cat. No. TH269517: 6G9, Thermo Fischer Scientific, Rockford, USA) or mouse anti-GAD67 (1:1000, cat. No ab26116, Abcam, Cambridge, GBR). Next day, slices were washed in PBS (6x 5-minute washes) and incubated for 2 hours at room temperature in blocking buffer containing the polyclonal secondary antibody goat anti-mouse IgG conjugated with Alexa Fluor^TM^ 594 (1:1000, cat. No. A11005, Life Technologies Co., Eugene, USA). After washing in PBS (6x 5-minute washes) the slices were mounted in adhesion microscope slides with Fluoroshield™ mounting media. Images of the dmPFC contralateral to the neuronal tracer microinjection site were captured at 25x, 200x and 400x magnification on an epifluorescence microscope (Axio Imager.D2) using the Zen Pro 2.0 software for the visualization of the neuronal tracer migration and quantification of CAMKII alpha staining, GAD67 staining and Fluoro-Gold^TM^/CAMKII alpha or Fluoro-Gold^TM^/GAD67 double staining.

#### 2.4.1. Immunofluorescence quantification

The background from the obtained images was manually subtracted based on the color histogram using the Zen Pro 2.0 software. The same criterion was applied to all images. The total fluorescence of the neuronal tracer Fluoro-Gold^TM^, CAMKII alpha, and GAD67 was quantified by the corrected total cell fluorescence (CTCF) method using the software ImageJ 1.54f (National Institutes of Health, Bethesda, USA). For that, the area and the integrated density of the dmPFC, and the integrated density of a background region of the slide were measured. Then, the final value was obtained according to the following formula: CTCF = integrated density of the slice – (area of the slice x integrated density of the background readings) (Canto-de-Souza et al., 2025). The total number of double-stained cells was manually counted using the software ImageJ 1.54f and the final value was corrected to the dmPFC area of each slice (Santos-Costa et al., 2021). For each measurement, values from 2 adjacent slices per mouse were obtained and averaged. All analyses were performed by an investigator blinded to the experimental groups.

### 2.5. Psychosocial stress

We used a modified version of the resident-intruder model (Pryce and Fuchs, 2017) and the witness social defeat stress (WSDS) paradigm (Iñiguez et al., 2018) as a protocol for the PSS, allowing simultaneous episodic exposure of both females and males to the stressor stimulus, as already performed in our laboratory (Canto-de-Souza et al., 2025).

Accordingly, male experimental subjects (intruders) were exposed to the home cage of an aggressive male conspecific (resident), while one or two female experimental subjects (witnesses) were watching the agonistic encounter between males to observe aggression toward the intruder and subsequent submission. To accommodate the witness in the resident’s home cage, an acrylic T-shaped removable perforated divider (207 x 216 x 100 mm, custom-made, Master One, Ribeirão Preto, BRA) was positioned on one side (the witnessing area), allowing two-thirds of the space for agonistic encounters. Witnessing mice had no visual contact with each other. Each PSS episode lasted 15 minutes, divided into three phases: first, the intruder (male experimental subject) was placed in a wire-mesh protective cage (155 x 106 x 48 mm, custom-made, Master One) for 5 minutes. The intruder was then removed from the protective cage and exposed to resident attacks for 5 minutes, while the witness observed the interaction. Finally, the intruder was returned to the protective cage for an additional 5 minutes, during which the witness remained in the resident’s home cage within the witnessing area. At the end of the PSS episode, intruders and witnesses were returned to their respective home cages. Each encounter was scheduled such that the resident was unfamiliar to both the intruder and the witnesses. In the repeated PSS (rPSS) animals were exposed to the protocol for 10 consecutive days, once daily. In the subthreshold PSS (sPSS), animals were exposed to a single encounter. As a control for the PSS, a non-aggressive interaction (NAI) and its observation were conducted. In NAI, two co-housed familiar males were placed in a novel cage to interact for 5 minutes, while females, confined to a protective area using a T-shaped removable perforated divider, witnessed the interaction. We used a shorter period of social interaction in NAI to avoid aggressiveness between the males. It was performed once a day for ten days (rNAI), as the control for the rPSS or as a single encounter (sNAI) as the control for the sPSS.

### 2.6. Physical state evaluation

Physical state was evaluated by analyzing the changes in body weight and coat state deterioration, as an indicator of the individual’s motivation for self-centered activities and apathy-like behavior (Planchez et al., 2019).

Before the first aggressive encounter in the PSS protocol (initial measurement) and the social interaction test (SIT, final measurement), experimental subjects were evaluated regarding body weight and the condition of their fur. Fur condition was evaluated using a scale from 3 (healthy and well-cared fur) to 0 (unhealthy and dirty fur, hair loss, and piloerection), and intermediate states were assigned using 0.5-point decrements. Coat state deterioration and body weight change were expressed as the difference between the initial and final measurements.

### 2.7. Social interaction test

Social avoidance was evaluated in female and male mice using the SIT (Morais-Silva et al., 2023), 24 hours after the last PSS episode. The SIT was performed in an acrylic open arena (420 x 420 x 250 mm, custom-made, Master One) containing a perforated acrylic contention box (100 x 100 x 240 mm, custom-made, Master One) centered on one wall. The arena was virtually divided into an interaction zone (150 mm width zone near the contention box) and two avoidance zones (150 x 150 mm corners in the opposite wall relative to the contention box).

The experimental animals were put in the center of the arena facing the empty contention box for 150 seconds for the evaluation of the basal exploratory activity (no-target session). Then, animals were removed from the arena, and a non-aggressive conspecific (not used in the experiments) was placed in the contention box. The experimental animals were put again in the center of the arena facing the contention box for an additional 150 seconds for the evaluation of the social approaching/avoidance (target session). Animals’ behavior was recorded for the evaluation of the time spent in the interaction and avoidance zones, and the total distance traveled in the apparatus using the software ANY-maze® (Stoelting Co., Wood Dale, USA). The apparatus was cleaned with a hydroalcoholic solution (20% v/v) between subjects, and experiments were performed under dim red light (5 lux in the center of the apparatus) during the light phase of the circadian cycle.

### 2.8. Elevated plus maze (EPM)

Anxiety-related behaviors were assessed in the EPM following the original description (Lister, 1987) and previous work from our lab (Canto-de-Souza et al., 2025; Morais-Silva et al., 2023), 24 hours after the SIT. The EPM consists of a custom-made wooden cross-shaped apparatus elevated 385 mm above the floor, with two open arms (no protective walls, 300 x 50 x 2.5 mm) and two closed arms (surrounded by transparent glass walls, 300 x 50 x 150 mm) connected by a central platform (50 mm x 50 mm).

During the test, animals were placed in the center of the apparatus facing one open arm for 5 minutes. The number of entries and time spent in the open and closed arms were recorded via a camera connected to a computer and analyzed using the software ANY-maze®. These data were used to calculate the percentage of open arm entries (number of open arm entries/total entries into open and closed arms x 100) and the percentage of time spent in open arms (time in open arms / 300 x 100) (Rodgers and Johnson, 1995). The apparatus was cleaned with a hydroalcoholic solution (20% v/v) between subjects, and experiments were conducted under dim lighting (50 lux at the center of the apparatus) during the light phase of the circadian cycle.

### 2.9. Experimental procedures

#### 2.9.1. Experiment 1: Chemogenetic activation of the neuronal projections from the left to the right dorsal medial prefrontal cortex during repeated psychosocial stress in female and male mice

We used a chemogenetic approach to activate the direct projections from the LdmPFC to the RdmPFC during exposure to the rPSS, aiming to evaluate if the activation of such projections could prevent stress effects in female and male mice. Thirty male and 30 female mice underwent the stereotaxic surgery for the microinjection of the pan-neuronal AAV5-Cre in the LdmPFC and the retrograde Cre-dependent excitatory DREADD rgAAV-DIO-hM3Dq in the RdmPFC. The retrograde Cre-dependent rgAAV-DIO-mCherry was used in the RdmPFC of the control groups. Fourteen days later, animals were submitted to the rPSS protocol. CNO was administered 30 minutes before each aggressive encounter (or rNAI in control groups). Physical state was evaluated before the first PSS episode and before the SIT. Twenty-four hours after the last aggressive encounter, animals were tested in the SIT, and, 24 hours later, in the EPM. The experimental procedure is depicted in Figure 1A. A representative schematic coronal section of the microinjection site is available in Figure 1C.

#### 2.9.2. Experiment 2: Chemogenetic inactivation of the neuronal projections from the left to the right dorsal medial prefrontal cortex during the subthreshold psychosocial stress in female and male mice

We used a chemogenetic approach to inhibit the direct projections from the LdmPFC to the RdmPFC during exposure to the sPSS, aiming to evaluate if the inhibition of such projections could increase stress vulnerability in female and male mice. Thirty male and 32 female mice underwent the stereotaxic surgery for the microinjection of the pan-neuronal AAV5-Cre in the LdmPFC and the retrograde Cre-dependent inhibitory DREADD rgAAV-DIO-hM4Di in the RdmPFC. The retrograde Cre-dependent rgAAV-DIO-mCherry was used in the RdmPFC of the control groups. Fourteen days later, animals were submitted to the sPSS protocol. CNO was administered 30 minutes before the aggressive encounter (or NAI in control groups). Twenty-four hours later, animals were tested in the SIT, and 24 hours later, in the EPM. The experimental procedure is depicted in Figure 1B. A representative schematic coronal section of the microinjection site is available in Figure 1C.

#### 2.9.3. Experiment 3: Characterization of the interhemispheric projections of the dorsal medial prefrontal cortex

To characterize the neuronal projections between the left and right hemispheres of the dmPFC, 10 male and 11 female mice underwent the stereotaxic surgery for the microinjection of the retrograde neuronal tracer (Fluoro-Gold™) into the LdmPFC or RdmPFC, as described above. Twenty-one days after the microinjection, animals were euthanized, and the dmPFC was processed for the quantification of Fluoro-Gold™, CAMKII alpha, and GAD67 markers, as previously described. A representative schematic coronal section of the microinjection site is available in Supplementary Material S2 (Figure S2A).

### 2.10. Statistics

Data were expressed as the mean ± SEM. Statistics were performed using the software Statistica 14 (TIBCO Software Inc., Palo Alto, USA), and graphs were generated in the GraphPad Prism 8 software (GraphPad Software Inc., Boston, USA). Results were analyzed by two-way or repeated measures Analysis of Variance (ANOVA), according to the specificity of each dataset, considering the factors PSS (rNAI or rPSS; sNAI or sPSS), AAV [mCherry or DREADD (hM3Dq or hM4Di)], the repeated factor session (no target or target), sex (female or male), and hemisphere (LdmPFC or RdmPFC). In cases where ANOVA showed significant differences (p ≤ 0.05), the Newman-Keuls post hoc test was performed. Except for experiment 3, the female and male datasets were analyzed separately.

## 3. Results

### 3.1. Chemogenetic activation of the neuronal projections from the left to the right dorsal medial prefrontal cortex prevents coat state deterioration induced by repeated psychosocial stress in female and male mice

The rPSS induced a deterioration of the fur quality in both male and female mice, while hM3Dq activation in the LdmPFC projections to the RdmPFC prevented rPSS-induced coat state deterioration (Figure 2A and 2C). The two-way ANOVA of the coat state deterioration in males showed a significant effect for the factor rPSS (F_1,26_ = 28.87; p < 0.001), AAV (F_1,26_ = 5.66; p < 0.05), and for the interaction between rPSS and AAV (F_1,26_ = 9.20; p < 0.01). The Newman Keuls post hoc test revealed that the rPSS/mCherry group presented an increase in coat state deterioration, relative to rNAI/mCherry (p < 0.001), rNAI/hM3Dq (p < 0.001), and rPSS/hM3Dq (p < 0.05) groups (Figure 2A). In females, the two-way ANOVA of the coat state deterioration showed a significant effect for the interaction between rPSS and AAV (F_1,26_ = 7.76; p < 0.01), while the post hoc test revealed a significantly increased coat state deterioration in the rPSS/mCherry group relative to the rNAI/mCHerry (p < 0.05) and rPSS/hM3Dq (p < 0.05) groups (Figure 2C).

The rPSS exposure decreased the body weight of both male and female mice (Figures 2B and 2D). On the other hand, the chemogenetic activation of the LdmPFC projections to the RdmPFC was not effective in preventing such alterations. In females, the repeated hM3Dq activation also induced a decrease in body weight. The two-way ANOVA of the body weight changes in males showed a significant effect for the factor rPSS (F_1,26_ = 7.69; p < 0.05), where rPSS-exposed males lost more weight relative to rNAI-exposed males (p < 0.05), independent of the viral vector administered (Figure 2B). The two-way ANOVA of the body weight changes in females showed a significant effect for the factors rPSS (F_1,26_ = 10.06; p < 0.01) and AAV (F_1,26_ = 5.64; p < 0.05), but not for the interaction between the factors. The rPSS-females lost more weight relative to rNAI-females (p < 0.01), regardless of hM3Dq activation, while the hM3Dq-females lost more weight relative to mCherry-females (p < 0.05), regardless of the rPSS exposure (Figure 2D).

Off-target animals in Experiment 1 were analyzed to exclude the influence of the viral vector itself on the behavioral responses to rPSS. The data is available as supplementary material (Supplementary Material 3). We find no effects of off-target viral vector expression in any of the parameters analyzed.

### 3.2. Chemogenetic activation of the neuronal projections from the left to the right dorsal medial prefrontal cortex prevents social avoidance, but not the decrease in locomotor activity, induced by repeated psychosocial stress in male mice

The exposure to the rPSS induced social avoidance and decreased exploratory activity in male but not female mice. On the other hand, the chemogenetic activation of the LdmPFC projections to the RdmPFC during the agonistic encounters prevented only the social withdrawal induced by rPSS in male mice, without altering the locomotor deficits (Figure 3).

The repeated measures ANOVA of the time spent in the interaction zone during the SIT in males showed a significant effect for AAV (F_1,26_ = 4.45; p < 0.05), session (F_1,26_ = 40.45; p < 0.001), and for the interaction between rPSS and AAV (F_1,26_ = 16.95; p < 0.001), rPSS and session (F_1,26_ = 25.65; p < 0.001), and rPSS, AAV, and session (F_1,26_ = 12.01; p < 0.01). The Newman-Keuls post hoc test revealed a significant decrease in the time spent in the interaction zone when the social target was present in the rPSS/mCherry group, relative to time spent in the interaction zone during target session in rNAI/mcherry (p < 0.001), rNAI/hM3Dq (p < 0.001), and rPSS/hM3Dq (p < 0.001) groups. Moreover, there was an increase in time spent in the interaction zone in the target session compared to no target session in rNAI/mCherry (p < 0.001), rNAI/hM3Dq (p < 0.01), and rPSS/hM3Dq groups (p < 0.05) (Figure 3A).

For the time spent in the avoidance zone in males, the repeated measures ANOVA showed a significant effect for the interaction between rPSS and session factors (F_1,26_ = 11.83; p < 0.01). The post hoc test revealed a decrease in time spent in the avoidance zone when the social target was present in rNAI-exposed animals (p < 0.05), regardless of hM3Dq activation. Additionally, rPSS-exposed animals showed an increase in time spent in the avoidance zone in the target session relative to rNAI-exposed animals (p < 0.05), regardless of hM3Dq activation (Figure 3B).

There was a significant effect for rPSS (F_1,26_ = 15.99; p < 0.001) and session factors (F_1,26_ = 97.68; p < 0.001) on the total locomotor activity in the SIT in male mice according to the repeated measures ANOVA. There was a decrease in the total distance traveled in the apparatus in the target session compared to the no target session (p < 0.001), while the rPSS-exposed animals showed a significant decrease in the total distance traveled regardless of the hM3Dq activation or session (p < 0.001) (Figure 3C).

In females, the repeated measures ANOVA of the time spent in the interaction zone showed a significant effect for the session factor (F_1,26_ = 45.89; p < 0.001). Female mice spent more time in the interaction zone when the social target was present, regardless of the rPSS exposure or hM3Dq activation (p < 0.001) (Figure 3D). The repeated measures ANOVA showed a significant effect for the session factor (F_1,26_ = 10.91; p < 0.01) in the time spent in the avoidance zone in female mice. Female mice spent less time in the avoidance zone in the target session relative to the no target session, regardless of rPSS exposure or hM3Dq activation (p < 0.01) (Figure 3E). Similarly, the repeated measures ANOVA showed a significant effect for the session factor (F_1,26_ = 79.67; p < 0.001) on the total locomotor activity in the SIT in female mice. There was also a decrease in distance traveled in the apparatus in the target session relative to the no target session (p < 0.001), regardless of rPSS exposure or hM3Dq activation (Figure 3F).

### 3.3. Chemogenetic activation of the neuronal projections from the left to the right dorsal medial prefrontal cortex does not impact the behavioral alterations in the elevated plus maze induced by repeated psychosocial stress in female and male mice

The rPSS exposure affected male and female behavior in the EPM in a sex-specific manner, while the activation of the LdmPFC neuronal projections to the RdmPFC did not affect rPSS effects. In males, rPSS exposure for 10 days decreased the number of entries in the closed arms of the apparatus, while in females, rPSS exposure had an anxiogenic-like effect, decreasing the time spent in the open arms (Figure 4).

The two-way ANOVA of the number of closed arms entries in male mice showed a significant effect for the rPSS factor (F_1,24_ = 6.58; p < 0.05). The rPSS-exposed animals showed a reduction in the number of entries in the closed arms relative to rNAI-exposed animals (p < 0.05), regardless of hM3Dq activation (Figure 4A). The two-way ANOVA did not show any significant effect on the percentage of entries in the open arms (Figure 4B) or the percentage of time spent in the open arms (Figure 4C) in male mice.

There were no significant effects in the number of closed arms entries (Figure 4D) or percentage of open arms entries (Figure 4E) in female mice. On the other hand, there was a significant effect for the rPSS factor (F_1,26_ = 4.90; p < 0.05) in the percentage of time spent in the open arms in female mice. The rPSS-exposed females showed a decrease in time spent in the open arms compared to rNAI-exposed females (p < 0.05), regardless of hM3Dq activation (Figure 4F).

### 3.4. Chemogenetic inhibition of the neuronal projections from the left to the right dorsal medial prefrontal cortex increases the vulnerability to social avoidance and locomotor alterations induced by psychosocial stress in male mice

As expected, sPSS exposure did not affect male mice’s behavior in the SIT. On the other hand, the chemogenetic inhibition of the LdmPFC projections to the RdmPFC increased the vulnerability of male mice to sPSS, as it decreased social interaction and induced locomotor alterations in the EPM. In female mice, sPSS exposure increased social approach, as female mice that witnessed one episode of social defeat increased their time spent in the target zone when the social target was present. The hM4Di activation in female mice increased the distance traveled in the apparatus when a social target was absent (Figure 5).

The repeated measures ANOVA showed a significant effect for the factors sPSS (F_1,26_ = 5.83; p < 0.05), AAV (F_1,26_ = 14.24; p < 0.001), session (F_1,26_ = 48.77; p < 0.001), and for the interaction between sPSS and session (F_1,26_ = 10.65; p < 0.01) and sPSS, AAV, and session (F_1,26_ = 5.10; p < 0.05) for the time spent in the interaction zone in male mice. The Newman Keuls post hoc test revealed a significant increase in time spent in the interaction zone in the target session relative to no target session in sNAI/mCherry (p < 0.001), sNAI/hM4Di (p < 0.001), and sPSS/mCherry (p < 0.001) groups, but not in the sPSS/hM4Di group. Moreover, the sPSS/hM4Di group spent less time in the interaction zone when the social target was present compared to sNAI/mCherry (p < 0.001), sNAI/hM4Di (p < 0.001), and sPSS/mCherry (p < 0.001) groups (Figure 5A). There were no significant effects in the time spent in the avoidance zone in male mice (Figure 5B).

There was a significant effect for the factor session (F_1,26_ = 173.94; p < 0.001) and for the interaction between sPSS and session (F_1,26_ = 8.67; p < 0.01) and sPSS, AAV, and session (F_1,26_ = 4.45; p < 0.05) on the distance traveled in the SIT apparatus in male mice. The Newman Keuls post hoc test revealed a significant decrease in the total distance traveled during the target session in all groups (p < 0.001). Moreover, the sPSS/hM4Di group presented a further decrease in locomotor activity during the target session relative to sNAI/mCherry (p < 0.01) and sNAI/hM4Di (p < 0.01) groups (Figure 5C).

The repeated measures ANOVA showed a significant effect for the sPSS factor (F_1,28_ = 5.85; p < 0.05), session factor (F_1,28_ = 37.04; p < 0.001), and for the interaction between sPSS and session (F_1,28_ = 5.68; p < 0.05) for the time spent in the interaction zone in female mice. The post hoc test revealed a significant increase in the time spent in the interaction zone when the social target was present in all groups (p < 0.05). In female mice exposed to the sPSS, time spent in the interaction zone during the target session was further increased compared to sNAI female mice, regardless of hM4Di activation (p < 0.01) (Figure 5D). The repeated measures ANOVA showed a significant effect for the interaction between session and AAV factor (F_1,28_ = 5.37; p < 0.05) for the time spent in the avoidance zone. However, the Newman-Keuls post hoc test did not reveal any significant differences (Figure 5E).

Concerning the total locomotor activity in female mice during the SIT, the repeated measures ANOVA showed a significant effect for the session factor (F_1,28_ = 153.42; p < 0.001) and for the interaction between sPSS and session (F_1,28_ = 5.18; p < 0.05) and AAV and session (F_1,28_ = 9.85; p < 0.01). The post hoc test revealed that both sNAI and sPSS-exposed females showed a decrease in total distance traveled in the target session compared to the no target session (p < 0.001). Moreover, hMD4i activation increased total locomotor activity in female mice during the no target session (p = 0.05), regardless of sPSS exposure (Figure 5F).

### 3.5. Chemogenetic inhibition of the neuronal projections from the left to the right dorsal medial prefrontal cortex increases anxiety-like behavior in male but not female mice in the elevated plus maze

The sPSS did not affect female or male mice’s behavior in the EPM. However, the inhibition of the neuronal projections from the LdmPFC to the RdmPFC increased anxiety-like behaviors in the EPM 24 hours later in male but not female mice (Figure 6).

There were no significant effects for the number of closed arm entries (Figure 6A) or the percentage of entries in the open arms (Figure 6B) in the EPM in male mice. On the other hand, the two-way ANOVA revealed a significant effect for the AAV factor (F_1,26_ = 3.89; p = 0.05) for the percentage of time spent in the open arms. The hM4Di-injected animals showed a significant decrease in the percentage of time spent in the open arms of the apparatus relative to mCherry-injected animals (p = 0.05) (Figure 6C).

The two-way ANOVA did not show any significant effect for the number of closed arms entries (Figure 6D), percentage of entries in the open arms (Figure 6E), or percentage of time spent in the open arms (Figure 6F) in the EPM in female mice.

### 3.6. The left dorsomedial prefrontal cortex exhibits a greater number of glutamatergic projections to the contralateral hemisphere compared to the right dorsomedial prefrontal cortex in mice

Surprisingly, the double staining of Fluoro-Gold^TM^ + CAMKII-alpha and Fluoro-Gold^TM^ + GAD67 markers revealed that both glutamatergic and GABAergic neurons project from one hemisphere of the dmPFC to another (Figure S2). A representative schematic coronal section of the microinjection site is available in Figure S2A. Figures S2B and S2C are representative photomicrographs of the microinjection site in the LdmPFC and RdmPFC, respectively. The photomicrographs in 400x magnification revealed the presence of both Fluoro-Gold^TM^ + CAMKII-alpha and Fluoro-Gold^TM^ + GAD67 double-labeled neurons in the LdmPFC (Figure S2D and S2E, respectively) and RdmPFC (Figure S2F and S2G, respectively).

Next, we quantified the number of double-labeled neurons in the LdmPFC and RdmPFC of female and male mice (Figure 7). The repeated measures ANOVA showed a significant effect for the hemisphere factor (F_1,17_ = 13.80; p < 0.01) on the number of interhemispheric glutamatergic projections. The Newman-Keuls post hoc test revealed a higher number of glutamatergic projections in the left hemisphere compared to the right (p < 0.01), regardless of sex (Figure 7A). The repeated measures ANOVA did not show any significant effects on the number of GABAergic projections (Figure 7B). Thus, our results suggest that the glutamatergic projections from the LdmPFC to the contralateral hemisphere are heavier relative to the RdmPFC contralateral projections.

The repeated measures ANOVA did not show any significant effects on the corrected total fluorescence of Fluoro-Gold^TM^, CAMKII-alpha, and GAD67, suggesting that there are no significant differences relative to the total number of projections (Fluoro-Gold^TM^ labeled neurons), total number of glutamatergic neurons (CAMKII-alpha labeled neurons) or total number of GABAergic neurons (GAD67 labeled neurons) (Figure 7C). Representative photomicrographs in 200x magnification are available in Figures 7D and 7E.

## 4. Discussion

Here we showed that the bidirectional chemogenetic modulation of the LdmPFC neuronal projections activity to the contralateral dmPFC during stress modulates the individual vulnerability to PSS in a sex-specific manner in mice. Additionally, there is a higher number of glutamatergic projecting neurons from the LdmPFC to the RdmPFC, which should be related to the behavioral outcomes found in our study.

Since the proposal of the hypothesis that a dysregulated functional lateralization of the mPFC may be a key factor in the development of stress-related disorders (Cerqueira et al., 2008), subsequent studies highlighted that the disruption of the tonic inhibitory activity from the LdmPFC over the RdmPFC is central in chronic stress pathophysiology. Our results indicate, for the first time, that the apathy-like effects in female and male mice and the social deficits in male mice induced by a chronic psychosocial stressor can be rescued by an increase in the activity of the direct neuronal projections from the LdmPFC to the contralateral dmPFC during PSS exposure, while the inhibition of such projections during a single stress episode increases the male mice vulnerability to PSS-induced social deficits and anxiogenic effects.

Based on our results, we can hypothesize that the maintenance of tonic activity of the LdmPFC projections to the contralateral dmPFC during chronic stress exposure can prevent stress induced RdmPFC overactivity and consequent social deficits. Indeed, others have shown that increased RmPFC activity is related to CSDS-induced social deficits (Faria et al., 2020; Lee et al., 2015) and to the anxiogenic effects of the LdmPFC inhibition in male mice (Costa et al., 2016; Santos-Costa et al., 2021), that can be prevented by LmPFC stimulation (Lee et al., 2015) or RdmPFC glutamatergic blockade (Santos-Costa et al., 2021). On the other hand, inhibiting the abovementioned projections during a single PSS episode may disrupt RdmPFC tonic inhibition by the LdmPFC, leading to an increased vulnerability to stress effects, at least in males. In this context, LdmPFC inhibition using cobalt chloride prolongs the anxiogenic-like effects of a single SDS exposure (Santos-Costa et al., 2021; Victoriano et al., 2020) and increases the activation of glutamatergic neurons in the RdmPFC in male mice (Santos-Costa et al., 2021).

A dysregulation of LdmPFC to RdmPFC projections activity seems to be related to rPSS-induced apathy-like effects in female and male mice and social deficits in male mice, but not to rPSS-related behavioral alterations in the EPM. However, disrupting LdmPFC activity during stressful situations seems to be enough to impair mPFC functional lateralization and increase anxiety-like behaviors and vulnerability to stress-induced social deficits, at least in male mice. It suggests that diverse neuronal projections from the LdmPFC are involved in distinct domains of stress-induced behavioral alterations and disruption of dmPFC functional asymmetries. In this sense, a significant proportion of neurons projecting to the contralateral mPFC often extend collateral projections to the amygdaloid complex (Cassell et al., 1989). Long-range excitatory projecting neurons from the contralateral mPFC have a biased preference, in layer 5 (L5) of the mPFC, for neurons targeting subcortical regions over neurons that project to cortical areas (Anastasiades and Carter, 2021). Thus, our results indicate that inhibiting the activity of the projecting neurons from the LdmPFC to RdmPFC is sufficient to induce an overall disruption of RdmPFC activity while enhancing the activity of the same neurons only rescues the stress-induced behavioral deficits controlled by this specific pathway.

Our findings revealed asymmetric glutamatergic connectivity between the hemispheres of the dmPFC, with LdmPFC to RdmPFC glutamatergic neurons outnumbering RdmPFC to LdmPFC projections. Thus, the proposed tonic inhibition of the LdmPFC to the right hemisphere (Cerqueira et al., 2008) seems to be mediated by increased activation of the local RdmPFC GABAergic interneurons, which theoretically are activated by the glutamatergic signals originating from the LdmPFC. In this sense, there is an asymmetry between LdmPFC and RdmPFC electrical responses to the contralateral stimulation that suggest differences in excitation/inhibition balance between the dmPFC hemispheres, with the RdmPFC showing an increase towards excitatory activity (Pérez et al., 1990). The RdmPFC showed increased resistance to fatigability in response to repetitive LdmPFC stimuli and increased amplitude changes in response to double stimulation of the contralateral hemisphere (Pérez et al., 1990). The increase in anxiety-like behaviors in the EPM in male mice or the prolongation of SDS-induced anxiogenesis in male mice related to LdmPFC inhibition is dependent on NMDA receptor activation in the RdmPFC (Santos-Costa et al., 2021; Victoriano et al., 2020), thus suggesting an increase in excitatory neurotransmission in the RdmPFC from other neuronal sources after LdmPFC blockade.

Additionally, we find both glutamatergic and GABAergic interhemispheric projecting neurons in the dmPFC. It was surprising since the glutamatergic pyramidal neurons constitute the majority of mPFC neurons and typically project to other regions, establishing long-range communication and serving as the primary source of informational output (Feldmeyer, 2015). In contrast, GABAergic neurons are primarily local interneurons, modulating both efferent glutamatergic projections and incoming signals (Staiger, 2015). However, there is evidence of mPFC GABAergic neurons forming long-distance projections, implicated in the modulation of aversive behavior, impulsivity, and learning (Cho et al., 2023; Lee et al., 2014; Saffari et al., 2016; Utashiro et al., 2024).

The higher number of glutamatergic projections from the LdmPFC to the RdmPFC does not exclude the participation of the GABAergic projecting neurons from the LdmPFC on the dmPFC functional asymmetries. They can act through the direct inhibition of the contralateral pyramidal neurons, increasing the tonic inhibitory activity exerted by the LdmPFC, or as modulators of the local RdmPFC interneuron activation by the contralateral glutamatergic projections. Although the literature lacks a characterization of such neurons, there is a study characterizing a class of long-range GABAergic projections within the dmPFC, which projects from the PrL to the Cg1/2. Those neurons are synaptically connected to GABAergic interneurons specifically on layer 1 (L1) of the Cg1/2, where they act through inhibition. The Cg1/2 L1 interneurons, in their turn, inhibit L5 pyramidal neurons. Thus, the activation of the PrL long-range GABAergic projections to the Cg1/2 results in an indirect inhibition of the Cg1/2 pyramidal cells (Utashiro et al., 2024). More studies are necessary to properly characterize the interhemispheric GABAergic projections and their role in dmPFC functional asymmetry.

It is important to disclose that using the absolute specificity of CaMKII-alpha expression for glutamatergic neurons should be carefully considered. Despite older evidence pointing towards a lack of CAMKII-alpha expression on cortical GABAergic interneurons (Wang et al., 2013), recent evidence suggests a functionally relevant expression of CAMKII-alpha on GABAergic interneurons, including in the mPFC (Veres et al., 2023). Still, studies have been demonstrating that CAMKII-alpha is highly enriched in glutamatergic pyramidal neurons (Basu et al., 2008; Egashira et al., 2018; Scheyltjens et al., 2015). Zhang et al. (2020) demonstrated that most CAMKII-positive neurons, identified by the administration of the viral vector AAV5-CaMKII-hM4Di-mCherry into wild-type mice, also express vGLUT1 mRNA (Zhang et al., 2020). Their chemogenetic inhibition attenuated cocaine-conditioned place preference acquisition and expression, while the chemogenetic activation of GAD67-positive neurons had no effect, suggesting that a substantial number of CAMKII-positive neurons are glutamatergic and functionally diverse from the GABAergic population. Additionally, the glutamic acid decarboxylase isoforms (GAD65 and GAD67) are enzymes responsible for GABA synthesis in the brain from glutamate, leading to the absence of glutamate release from GAD-positive neurons (Martin and Rimvall, 1993). In this sense, GABA is functionally released from GAD67-positive neurons (Chowdhury et al., 2019). Nevertheless, we believe that in our study, this impact is less relevant, and our dual-marker approach (CaMKII-alpha and GAD67) provides robust validation of neuronal phenotypes in the context of dmPFC interhemispheric-projecting neurons. Using this approach, we had a reliable answer regarding the GABAergic population, as the GAD67 marker seems to be specific for GABAergic cells. They accounted for a limited number of FluoroGold-positive cells, compared to the CaMKII-alpha/FluoroGold-positive cells. In this case, we can propose that our differences are related to the glutamatergic population since we did not find differences in the GAD67-positive cells.

The rPSS exposure did not induce social avoidance in female mice in the present study, but decreased self-care and increased anxiety-like behaviors as measured by the coat state deterioration and reduced open-arm exploration in the EPM, respectively. In C57BL/6 female mice subjected to 10 days of WSDS, stress exposure induced social avoidance in the SIT, increased depressive-like behaviors in the tail suspension test and sucrose preference test, and increased anxiety-like behaviors in the EPM (Iñiguez et al., 2018), along with decreased social preference in the 3-chamber social preference test (Morais-Silva et al., 2023). On the other hand, WSDS for 10 days did not affect the sociability of Swiss-Webster mice, but increased anxiety-like behaviors in the EPM and provoked memory impairment in the object recognition test (Canto-de-Souza et al., 2025). Whether differences in the stress protocol or mouse strain are involved in such discrepancies, or even whether female mice are less vulnerable to alterations in sociability after chronic stress exposure, remains to be determined.

The activation of LdmPFC projections to the contralateral hemisphere, shown in the present study, appears to have rescued the increase in apathy-like behaviors but not the increase in anxiety-like behaviors after rPSS exposure in female mice. Similarly, the inhibition of the above-mentioned projections did not increase the vulnerability of female mice to stress effects on the EPM after the sPSS protocol. Such results suggest a distinct role of the LdmPFC interhemispheric projections in female and male mice. In this sense, it is suggested that afferent projections to the mPFC are distinct between female and male rats (Laine et al., 2024). In addition, there are intrinsic differences between female and male mice on baseline glutamatergic neurotransmission within the mPFC (Knouse et al., 2022). The activation of neurons expressing vesicular glutamate transporter 3 from the median raphe nucleus to the dmPFC increases GABAergic neurotransmission and enhances fear extinction in female but not male mice (Collins et al., 2023). Moreover, mPFC responses to stress exposure seem to be distinct between female and male individuals. For instance, RRS exposure increases dendritic arborization in the mPFC of female rats but decreases in male rats (Garrett and Wellman, 2009). This increase in dendritic arborization in females is specific for neurons projecting to the basolateral amygdala (Shansky et al., 2010). In women, mPFC lesion increases the cortisol release in response to the trier social stress test (TSST), while it does not affect cortisol response in men. On the other hand, mPFC lesion increases heart rate and decreases heart rate variability during an orthostatic challenge in men but not women (Buchanan et al., 2010).

Another interesting sex-specific effect found in our study is the decrease in body weight due to the repeated chemogenetic activation of the neuronal projection from the LdmPFC to the RdmPFC in female mice. Evidence suggests that the mPFC plays a role in the neural control of feeding behavior and food valuation (Land et al., 2014). However, as far as we know, there are few studies looking for the role of the dmPFC on sex differences in feeding or motivation for food seeking. Three studies aimed for cFos expression on the dmPFC during food consumption (Greiner et al., 2024; Parsons et al., 2022; Reppucci and Petrovich, 2018), although no significant differences were found. On the other hand, a study showed that neuronal ensembles representing social and nonsocial rewards in the dmPFC are more distinct and less overlapping in female mice relative to male mice (Isaac et al., 2024). Thus, further studies are necessary to clarify the implications of the sex-specific effects related to the activation of the projections from the LdmPFC to RdmPFC.

The use of the EPM as the only behavioral test to evaluate anxiety-like behaviors limits a comprehensive assessment of the emotional state of mice following stress exposure. However, excessive exposure to behavioral tests could impact each other’s results (McIlwain et al., 2001; Shoji and Miyakawa, 2021). Thus, we delineated our protocol to keep the number of behavioral tests to a minimum, maintaining the evaluation of the behavioral domains that are impacted by chronic stress exposure and are related to psychiatric disorders (anxiety, motivation and social behavior) (American Psychiatric Association, 2013). We used the EPM since it is a classical behavioral test used for the evaluation of anxiety-like behaviors in rodents, and was extensively used in the studies that identified the implications of the mPFC functional asymmetry on stress (Costa et al., 2016; Santos-Costa et al., 2021; Sullivan and Gratton, 2002). The social interaction test was chosen due to its widespread use to identify the stress coping phenotypes after chronic stress exposure in rodents (Golden et al., 2011). Finally, the coat state was evaluated since it is used as an indicator of the individual’s motivation for self-centered activities and apathy-like behavior, and it is easily performed with minimum manipulation of experimental subjects (Planchez et al., 2019). Additionally, stress-induced alterations in the coat state and SIT are only reversed using chronic antidepressant treatments, reinforcing their predictive validity (Berton et al., 2006; Yalcin et al., 2008).

In conclusion, our results revealed an involvement of the monosynaptic projections from the LdmPFC to RdmPFC in the vulnerability to the behavioral alterations induced by PSS in female and male mice. We also showed a higher density of glutamatergic projections from the LdmPFC to the contralateral hemisphere than to the opposite. Those projections may be involved in the phenomenon of functional asymmetry of the dmPFC and its disruption by chronic stressors.

## Author contributions

GM-S and RLN-d-S conceived the idea and delineated the experiments. GM-S, BFF, and ILL performed experiments and analyses. GM-S wrote the main manuscript. RLN-d-S supervised the study. GM-S and RLN-d-S discussed and reviewed the results. GM-S, BFF, ILL, and RLN-d-S reviewed and edited the manuscript.

---

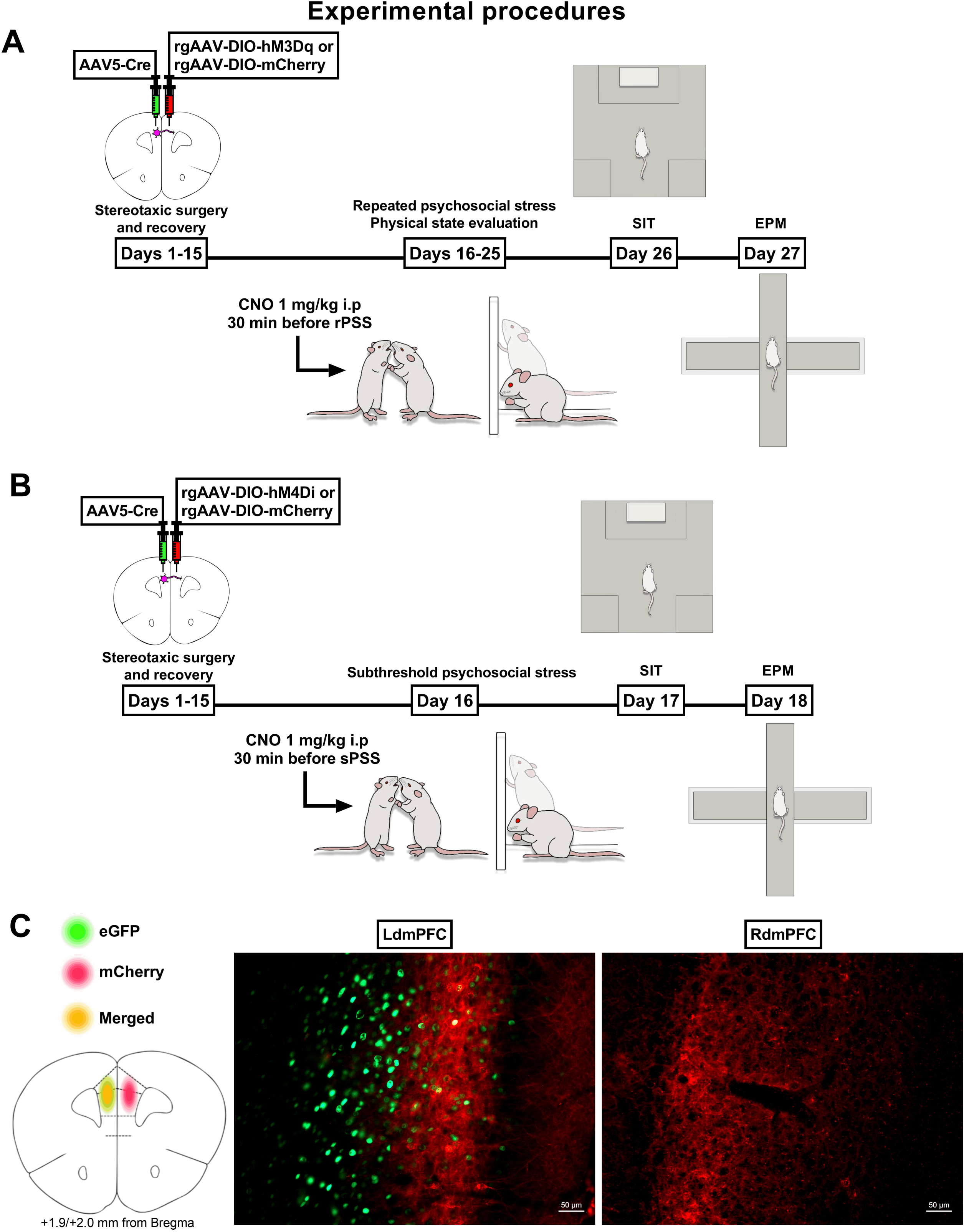

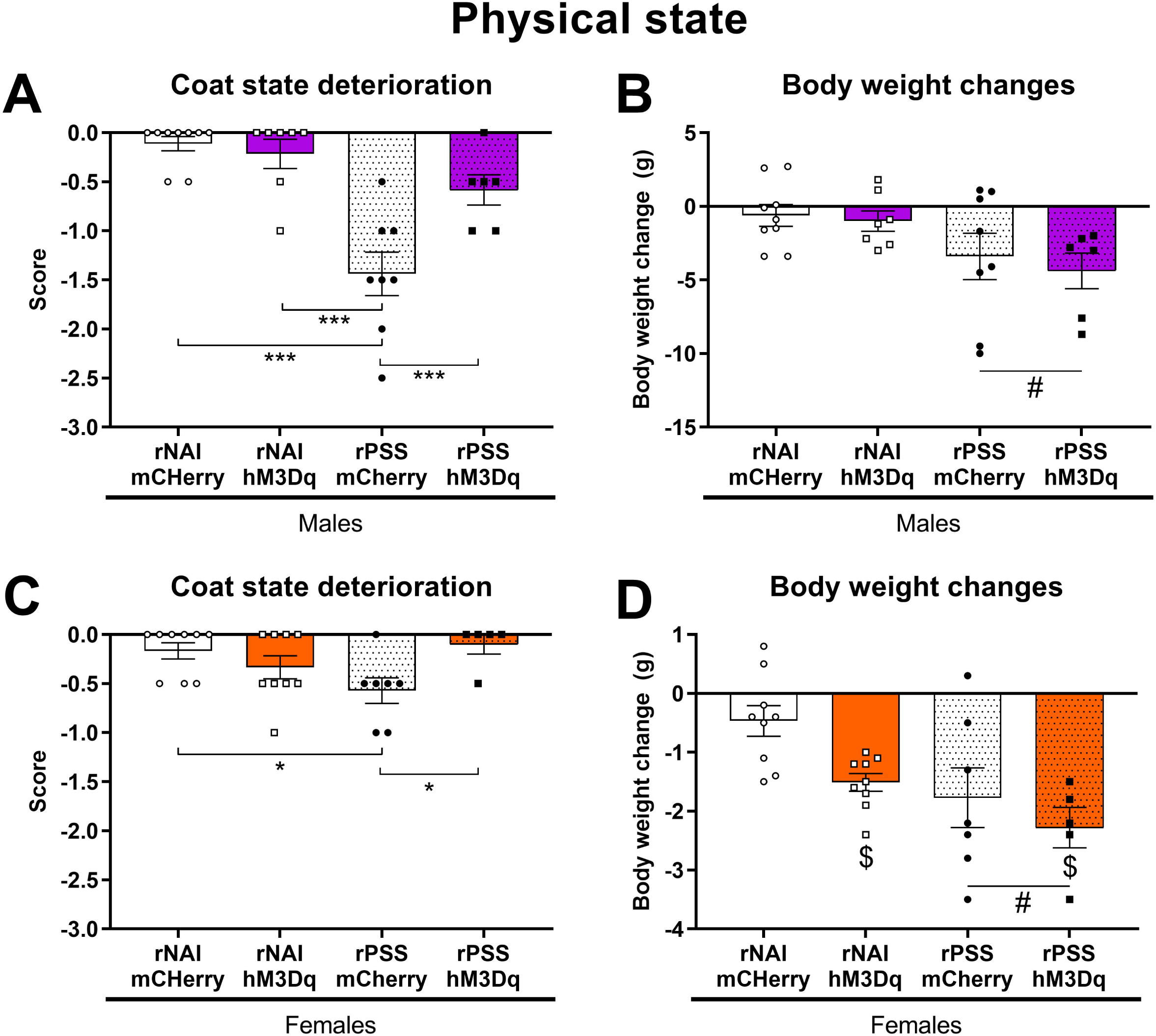

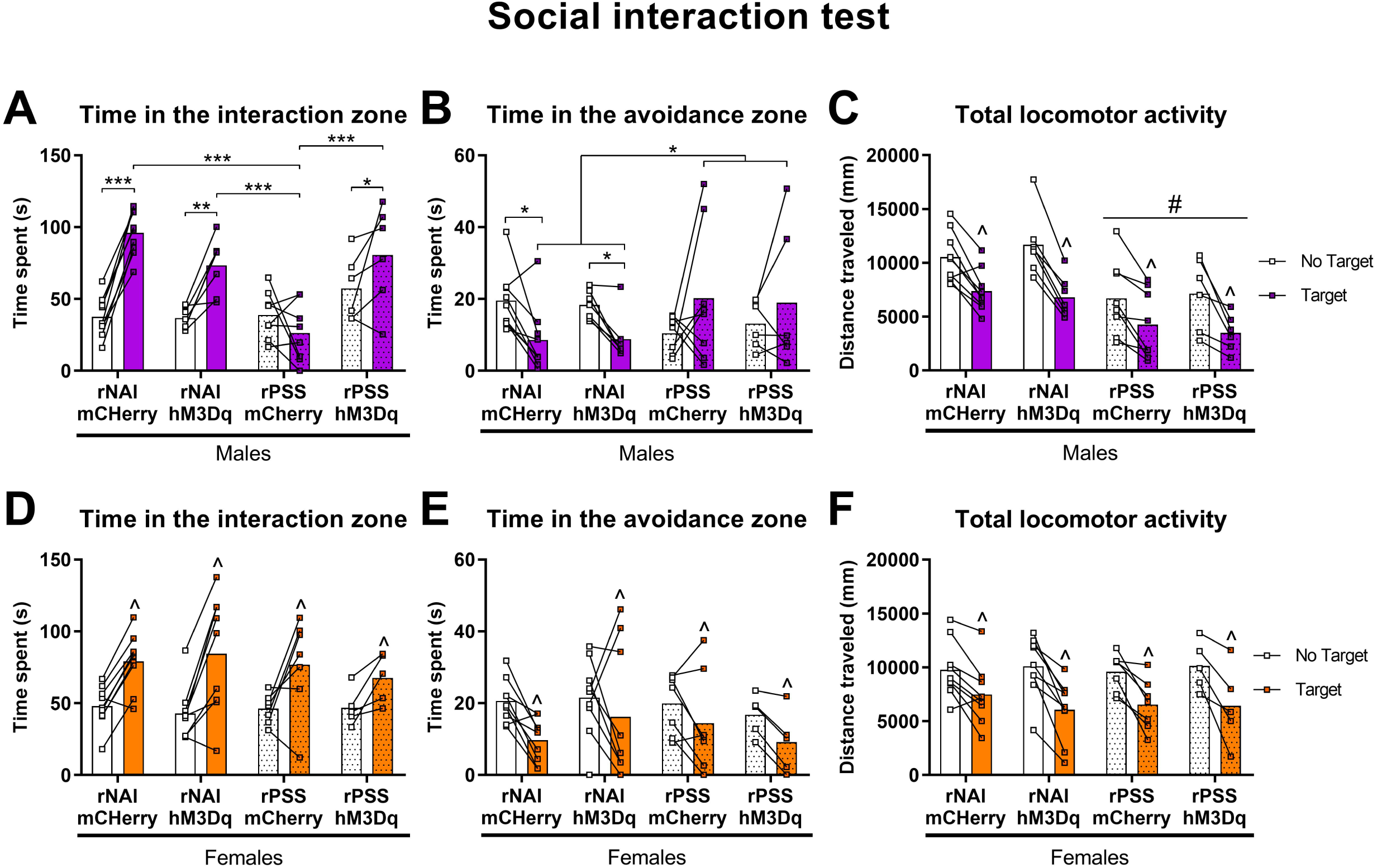

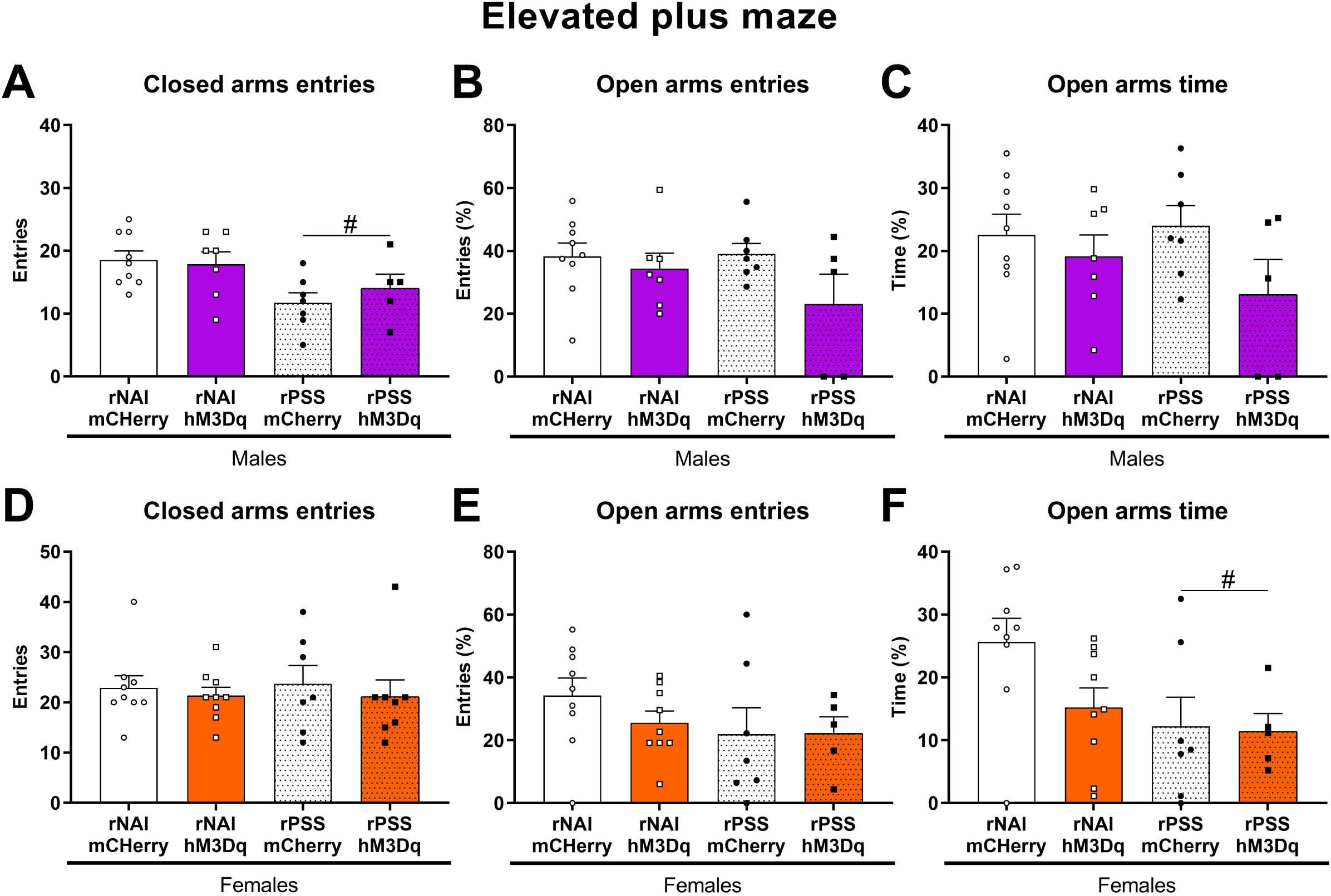

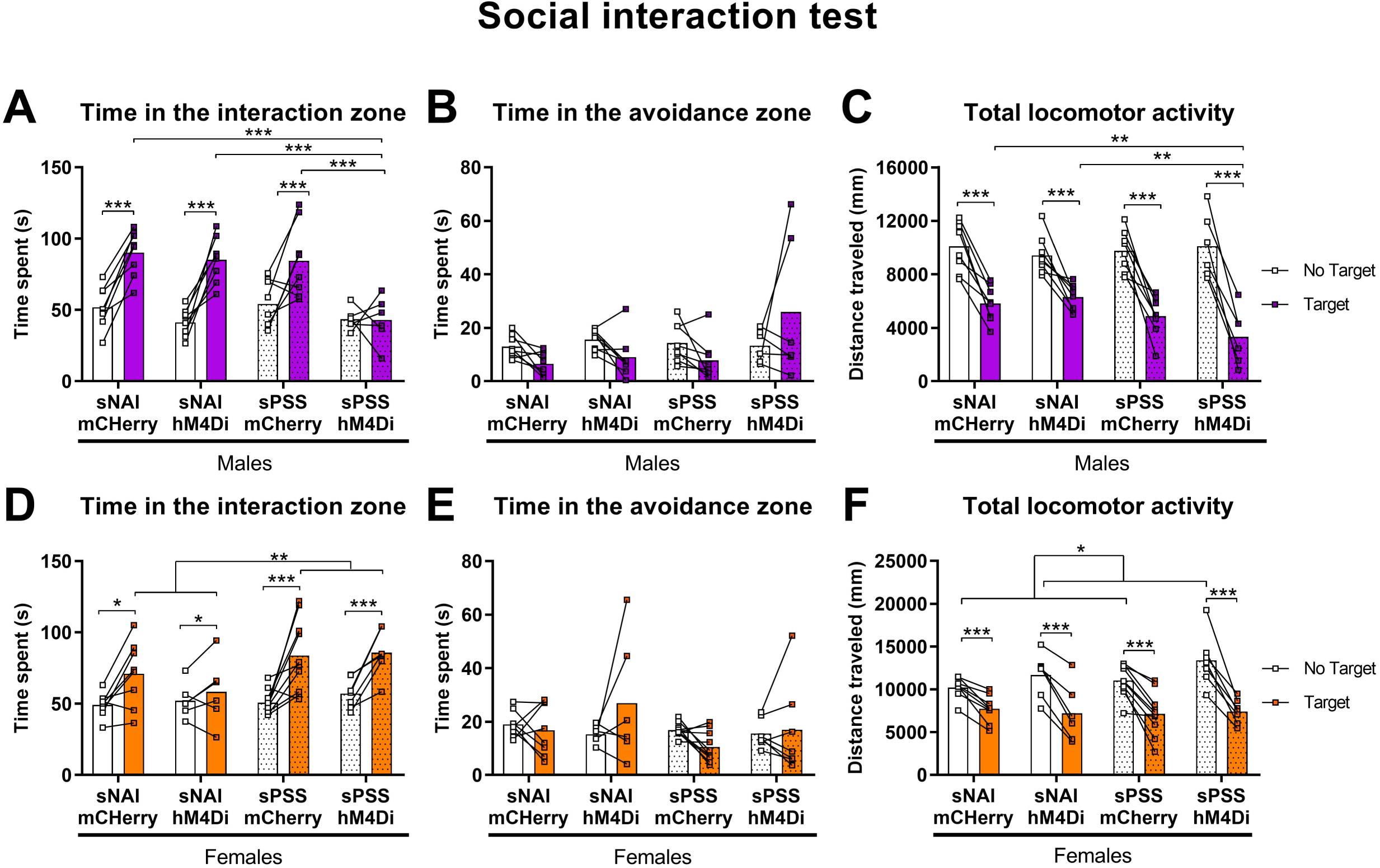

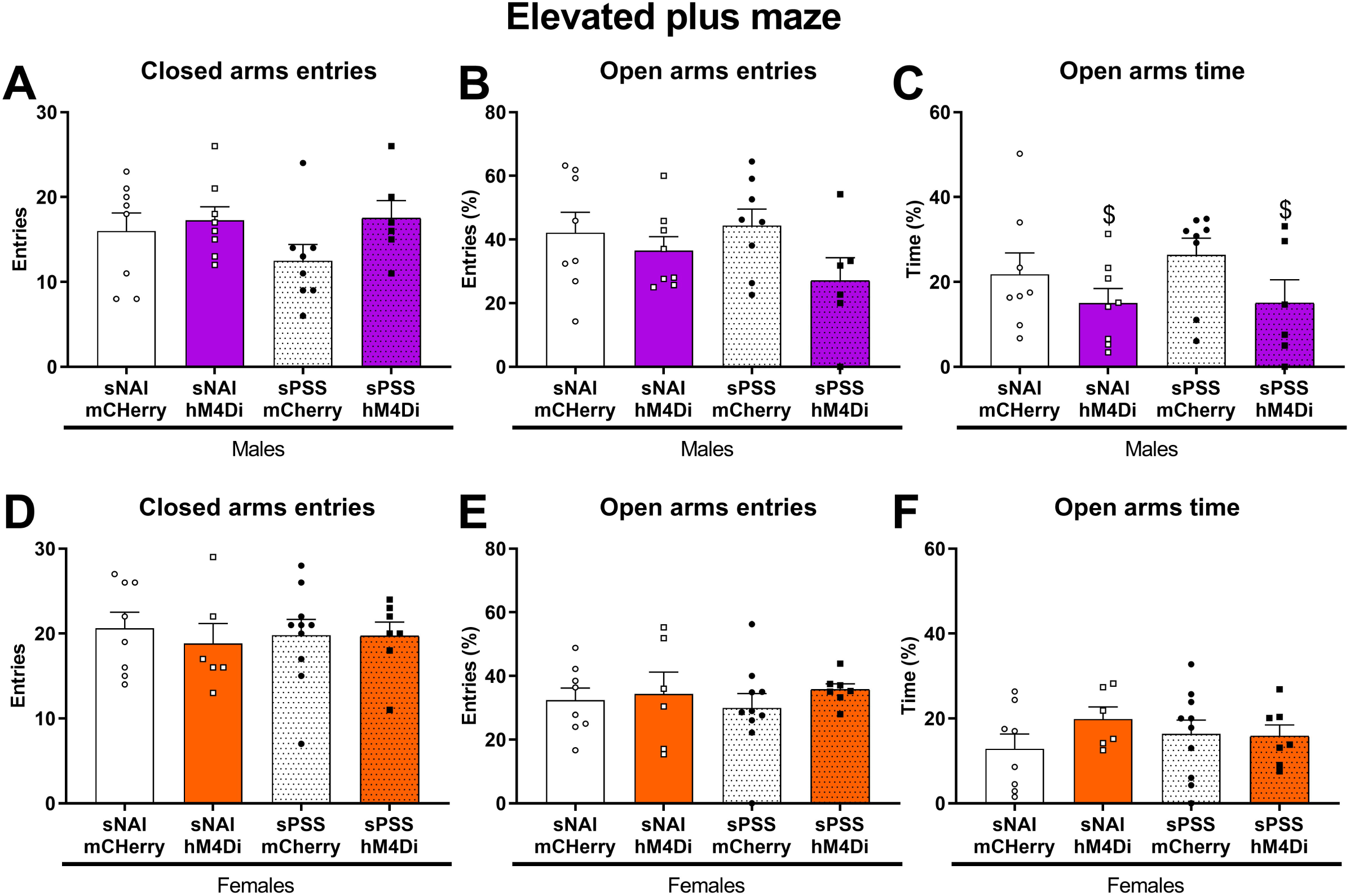

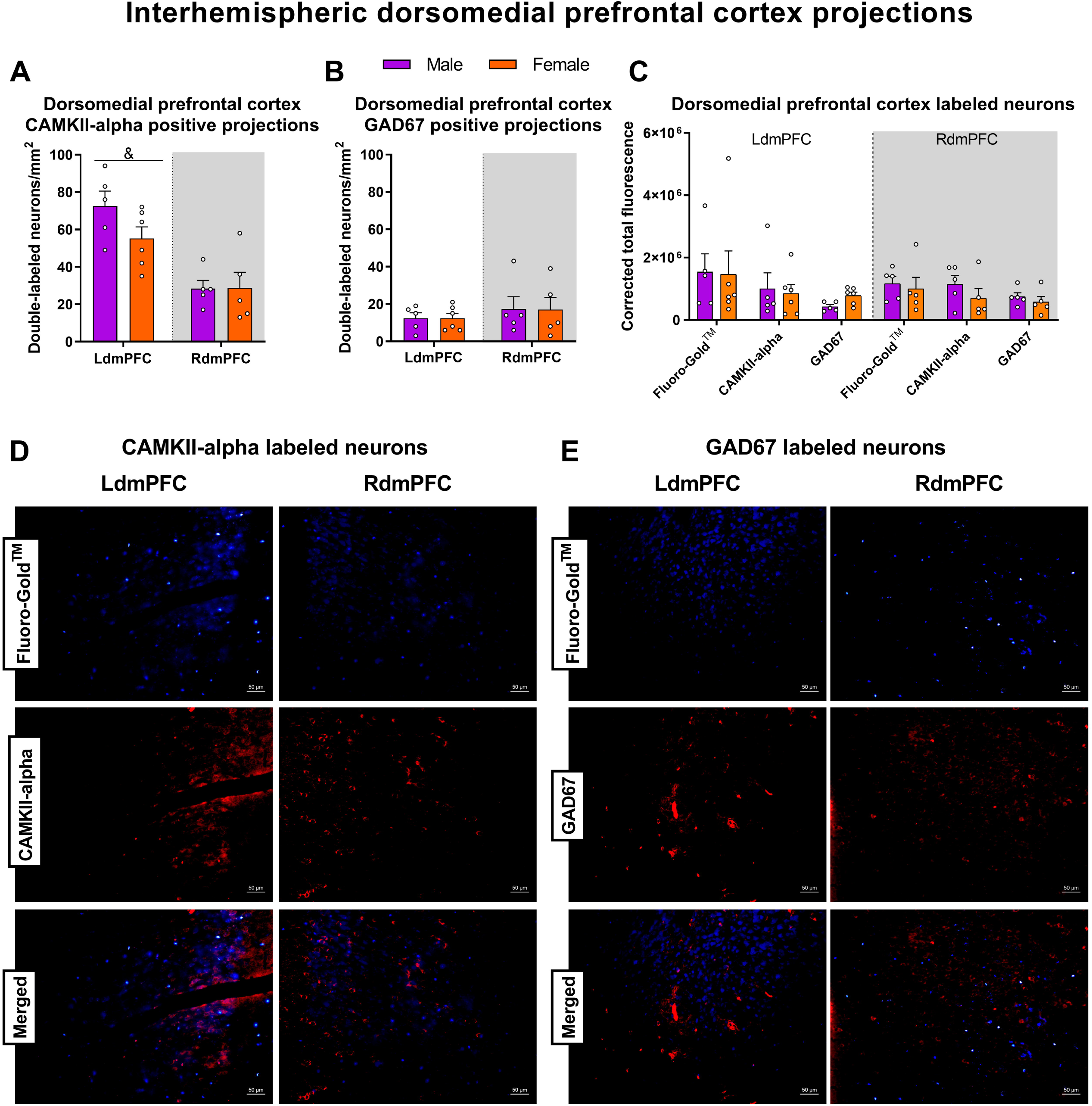

