## Supplementary material S1 for "Left-to-right dorsomedial prefrontal cortex interhemispheric projections mediate psychosocial stress vulnerability"

**Supplementary material S1 – Assessment of Clozapine-N-Oxide dose-dependent behavioral effects in the open field test in mice**

1. **Material and methods**
   1.1. *Subjects*

Thirty-nine Swiss-Webster mice (19 males and 20 females, 8 weeks old) were used. Animals were obtained from the Centro de Pesquisa e Produção de Animais (CPPA) at the São Paulo State University/UNESP (Botucatu, Brazil) and arrived at the local animal facility at 3 weeks old. Experimental subjects were grouped (age and sex-matched, 4-5 animals/cage) in individually ventilated polysulfone cages (240 x 251 x 386 mm, floor area 695 cm2, ref. 000121, ALBR Indústria e Comércio LTDA, Monte Mor, BRA) with sawdust bedding supplied with sterile cardboard tubes as environmental refinement (Ambient Trio®, Animal Pro, São Paulo, BRA). The facility was maintained under an artificial 12h light-dark cycle (lights on at 7 am), and controlled temperature (23 ± 1°C). Tap water (supplied by the distribution network) and food (Benelab, Qualy Nutrição Animal Indústria e Comércio Ltda., Lindóia, BRA) were freely available except during the brief periods of behavioral testing.

All procedures involving animals were previously approved by the local ethics committee for animal care and use (CEUA FCF/CAr protocol number 16/2021) and were conducted according to the principles of the Brazilian Guide for the Care and Use of Animals in Scientific Research (DBCA) issued by the National Council for Animal Experiments Control (CONCEA/MCTI). The DBCA was based on the NIH (National Research Council) Guide for the Care and Use of Laboratory Animals.

1.2. *Clozapine N-oxide*

Clozapine-N-oxide base (CNO) was obtained from the National Institute of Mental Health (NIMH) Chemical Synthesis and Drug Supply Program (catalog number C-929). It was dissolved in a vehicle solution containing dimethyl sulfoxide (DMSO) and 0.1M phosphate-buffered saline (PBS) in a 1:1 proportion (stock solution, 5 mg/mL) to prepare the following working solutions:

- CNO 0.1 mg/mL (final DMSO concentration 1%), administered in a volume of 0.1 mL/10g to achieve the dose of 1 mg/kg.
- CNO 0.5 mg/mL (final DMSO concentration 5%), administered in a volume of 0.2 mL/10g to achieve the dose of 10 mg/kg.

Vehicle-treated animals received a solution containing DMSO 5% in PBS in a volume of 0.2 mL/10g.

1.3. *Open field test*

The open field test (OFT) arena consists of an acrylic circular-shaped floor (300 mm diameter) surrounded by opaque walls (400 mm high), virtually divided into a central zone (150 mm diameter) and a peripheral zone. The animals were placed in the center of the arena and had their behavior recorded for 5 minutes for the evaluation of the total and central distance traveled, and the time spent in the center of the apparatus using the software ANY-maze® (Stoelting Co., Wood Dale, USA). OFT was performed 30 minutes after the administration of CNO (1 mg/kg or 10 mg/kg) or vehicle (n = 6-7 animals/group) intraperitoneally.

The apparatus was cleaned with a hydroalcoholic solution (20% v/v) between subjects, and experiments were performed under dim light (50 lux in the center of the apparatus) during the light phase of the circadian cycle.

1.4. *Statistics*

Data were expressed as the mean + SEM. Statistics were performed using the software Statistica 14 (TIBCO Software Inc., Palo Alto, USA), and graphs were generated in the GraphPad Prism 8 software (GraphPad Software Inc., Boston, USA). Results were analyzed by two-way Analysis of Variance (ANOVA), considering the factors treatment (vehicle, CNO 1 mg/kg or CNO 10 mg/kg), and sex (female or male). When ANOVA showed significant differences (p ≤ 0.05), the Newman-Keuls post hoc test was performed.

2. **Results**

The acute treatment with CNO at the dose of 10 mg/kg induced an anxiogenic-like effect in both female and male mice (Figure S1). On the other hand, CNO at the dose of 1 mg/kg did not induce behavioral alterations in the OFT.

The two-way ANOVA of the total distance traveled did not show any significant effect (Figure S1A). On the other hand, the two-way ANOVA of the distance traveled in the center showed a significant effect of the treatment factor (F_2,33_ = 5.41; p ≤ 0.01) (Figure S1B). There was a decrease in central distance traveled in CNO 10mg/kg treated animals relative to vehicle (p ≤ 0.05) and CNO 1 mg/kg groups (p ≤ 0.01), regardless of sex.

The two-way ANOVA of the time spent in the center showed a significant effect for the treatment factor (F_2,33_ = 9.03; p ≤ 0.001) (Figure S1C). CNO 10 mg/kg treated animals showed a significant decrease in time spent in the center of the apparatus relative to vehicle (p ≤ 0.001) and CNO 1 mg/kg groups (p ≤ 0.01), regardless of sex.

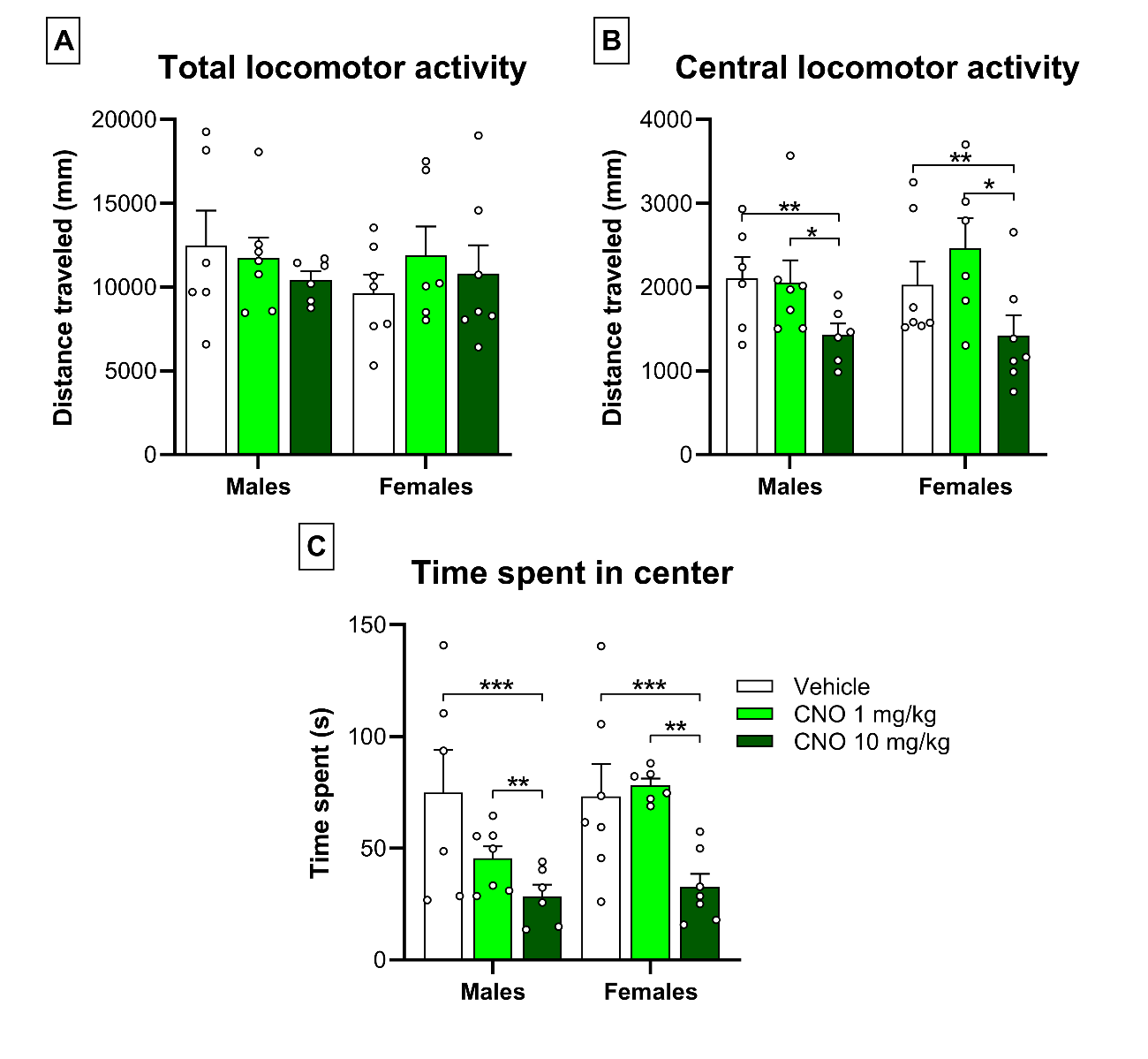

**Supplementary Figure S1.** Clozapine-N-Oxide (CNO) effects on the open field test (OFT). Male and female mice received CNO (1 mg/kg or 10 mg/kg) or vehicle 30 minutes before being tested in the OFT for 5 min. A, total distance traveled (in mm), B, distance traveled in the center (in mm) and C, time spent in the center (in sec). Data are presented as the mean (+ SEM) plus the individual values for each animal from male/vehicle (n = 6), male/CNO 1 mg/kg (n = 7), male/CNO 10 mg/kg (n = 6), female/vehicle (n = 7), female/CNO 1 mg/kg (n = 6) and female/CNO 10 mg/kg (n = 7) groups. *, **, ***, p ≤ 0.05, p ≤ 0.01, p ≤ 0.001 relative to the indicated group in the Newman-Keuls post hoc test.
