## Supplementary material S2 for "Left-to-right dorsomedial prefrontal cortex interhemispheric projections mediate psychosocial stress vulnerability"

**Supplementary material S2 - Phenotypic characterization of the interhemispheric dorsomedial prefrontal cortex (dmPFC) projections**

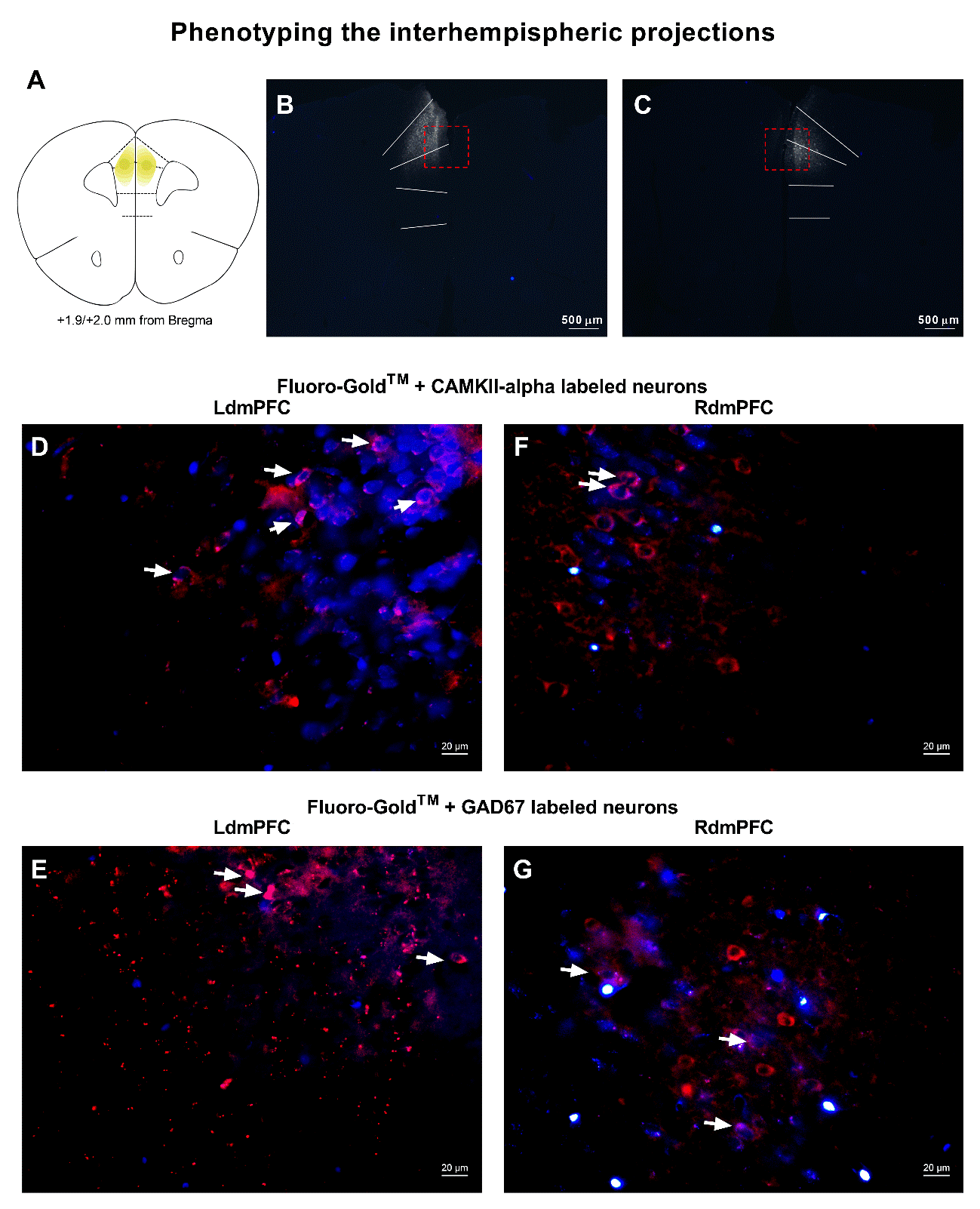

**Supplementary Figure S2.** Phenotypic characterization of the interhemispheric dorsomedial prefrontal cortex (dmPFC) projections in experiment 3. The animals received the retrograde neurotracer Fluoro-Gold^TM^ in the left (LdmPFC) or right (RdmPFC) hemisphere of the dmPFC and were euthanized 21 days later for the immunofluorescence. A, representative schematic coronal section of Fluoro-Gold^TM^ microinjection site in the LdmPFC or RdmPFC. Yellow ellipses represent the approximate spreading of the neurotracer fluorescence. B and C, representative low-magnification photomicrographs (25x) of the microinjection site and quantification site (dotted red square) when the neurotracer was administered in the LdmPFC or RdmPFC, respectively. Scale bar, 500 µm. D-G, representative high-magnification photomicrographs (400x) of Fluoro-Gold^TM^ + CAMKII-alpha or Fluoro-Gold^TM^ + GAD67 double-labeling in the LdmPFC or the RdmPFC contralateral to the microinjection site of the retrograde neurotracer. White arrows point to examples of double-labeled neurons. Scale bar, 20 µm.
