## Supplementary material S3 for "Left-to-right dorsomedial prefrontal cortex interhemispheric projections mediate psychosocial stress vulnerability"

**Supplementary material S3 – Analysis of off-target animals in the experiment “Chemogenetic activation of the neuronal projections from the left to the right dorsal medial prefrontal cortex prevents coat state deterioration induced by repeated psychosocial stress in female and male mice”**

The analysis of off-target animals did not reveal any significant effects of viral vector expression on the analyzed parameters.

The rPSS induced a deterioration of the fur quality in both male and female off-target mice (Figure S3A and S3B, respectively). The two-way ANOVA of the coat state deterioration in males showed a significant effect for the factor rPSS (F_1,26_ = 57.53; p < 0.001). In females, the two-way ANOVA of the coat state deterioration showed a significant effect for the rPSS factor (F_1,26_ = 6.91; p < 0.05).

The rPSS exposure induced a reduction in social interaction in male but nor female off-target mice in the SIT (Figure S3C and S3D, respectively). The repeated measures ANOVA of the time spent in the interaction zone during the SIT in males showed a significant effect for rPSS (F_1,26_ = 27.11; p < 0.001), session (F_1,26_ = 21.94; p < 0.001), and for the interaction between rPSS and session (F_1,26_ = 20.04; p < 0.001). The Newman-Keuls post hoc test revealed a significant decrease in the time spent in the interaction zone when the social target was present in the rPSS/mCherry and rPSS/hM3Dq groups. In females, the repeated measures ANOVA of the time spent in the interaction zone showed a significant effect for the session factor (F_1,26_ = 7.09; p < 0.05). Female mice spent more time in the interaction zone when the social target was present, regardless of the rPSS exposure or AAV administered.

Male, but not female off-target mice, exposed to the rPSS protocol showed a decrease in the number of entries in the closed arms of the EPM (Figure S3E and S3F, respectively). The two-way ANOVA of the number of closed arms entries in male mice showed a significant effect for the rPSS factor (F1,25 = 5.41; p < 0.05). The rPSS-exposed males showed a reduction in the number of entries in the closed arms relative to rNAI-exposed animals, regardless of AAV administered. There were no significant effects on the number of closed arms entries in female mice.

The two-way ANOVA showed no significant effects on the percentage of time spent in the open arms in male or female off-target mice (Figure S3G and S3H, respectively).

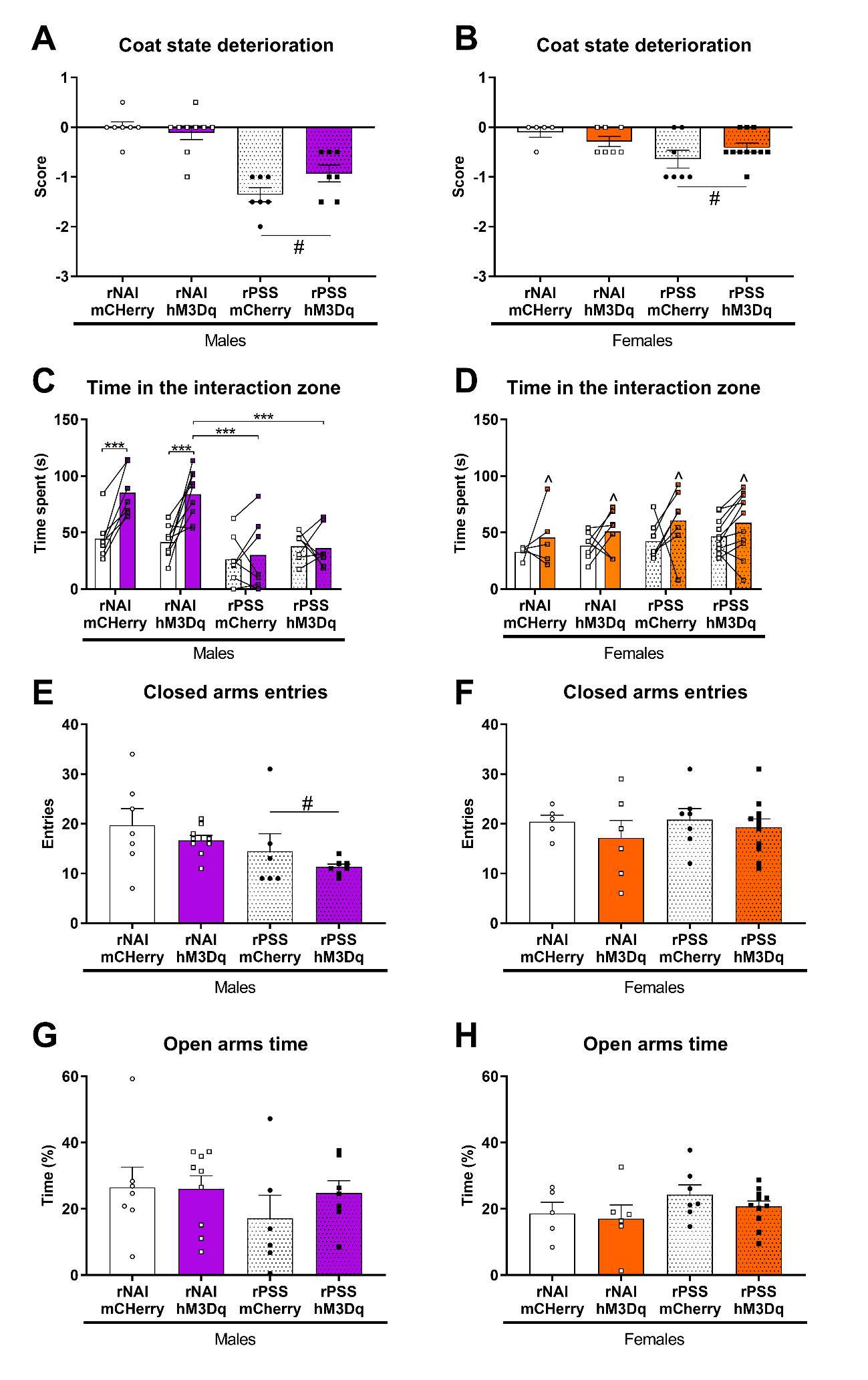

**Supplementary Figure S3.** Analysis of off-target animals in Experiment 1. A, B, coat state deterioration (score). C, D, time spent (in seconds) in the interaction zone during the no target and target sessions. E, F, closed arms entries. G, H, time spent (%) in the open arms of the EPM in male and female mice, respectively. Data are presented as the mean (+ SEM) plus the individual values for each animal from excluded animals of Male/rNAI/mCherry (n = 7), Male/rNAI/Hm3Dq (n = 9), Male/rPSS/mCherry (n = 7), Male/rPSS/hM3Dq (n = 7), Female/rNAI/mCherry (n = 5), Female/rNAI/Hm3Dq (n = 7), Female/rPSS/mCherry (n = 7), Female/rPSS/hM3Dq (n = 11) groups. *, **, ***, p ≤ 0.05, p ≤ 0.01, p ≤ 0.001 relative to the indicated group in the Newman-Keuls post hoc test. #, main effect (p ≤ 0.05) for the stress factor on the two-way ANOVA. ^, main effect (p ≤ 0.05) for the session factor on the repeated-measures ANOVA.
